# Proximate Composition and Mineral Analysis of Housefly Maggots and Mealworms Compared to Conventional Fish Meal and Soybean Meal

**DOI:** 10.1101/2025.05.18.654638

**Authors:** Gul E Nayab, Syed Basit Rashid, Sarzamin Khan, Asad Sultan, Irfat Ullah, Rafiullah Khan

**Author notes:** Corresponding author: Gul E Nayab.

## Abstract

The nutritional profiles of dried yellow mealworms and housefly maggots were analyzed and compared with conventional protein sources in poultry diets such as fish meal (FM) and soybean meal (SBM). Nutrient composition was determined through proximate analysis, while amino acid and mineral contents were analyzed using HPLC-based amino acid analyzer and atomic absorption spectrophotometer, respectively. Metabolizable energy (ME) was calculated to estimate energy content per kilogram of feed. The crude protein content of housefly maggots (57.8%) was comparable to that of fish meal (58.5%), which was the highest among the evaluated feeds, while mealworms had a slightly lower protein content (51.02%). The highest ether extract values were recorded for mealworms (27.3%) and maggots (26.9%), surpassing those in FM and SBM. The highest ME was calculated for housefly maggots (4,074 kcal/kg), followed by mealworms (3,988 kcal/kg) and FM (3,824 kcal/kg). Mineral profiling showed the highest calcium content in mealworms (5.23 g/kg), followed by maggots (5.13 g/kg). Phosphorus and zinc were recorded highest in housefly maggots, at 16.8 g/kg and 98.5 mg/kg, respectively. The limiting amino acids lysine, leucine, and isoleucine were highest in housefly maggots, followed by FM and mealworms, showing comparable values between the insect meals and FM. Methionine content was recorded highest in mealworms (2.3 g/100g), closely followed by housefly maggots (2.27 g/100g). The study demonstrated that housefly maggots and mealworms have a nutritive profile superior to soybean meal and comparable to fish meal.

## Introduction

Agriculture serves as the backbone of Pakistan’s economy, contributing significantly to the country’s GDP. One of the largest subsectors of agriculture is livestock [1], with poultry playing a key role in meeting the growing demand for protein. Poultry farming generates income and employment for numerous households, contributing a share of 1.4% to the total GDP [2]. Poultry farming in Pakistan has experienced rapid growth at a rate of 10-12% annually, driven by modern techniques, favorable climate, and supportive administrative regulations [3, 4]. The rapid growth of the poultry industry has brought challenges, particularly the rising costs of traditional protein sources such as soybean meal, fish meal, milk by-products, and processed animal proteins [5, 6]. This situation requires exploring alternative protein sources that can potentially reduce feed costs while maintaining or improving the performance and health of broiler birds.

Insects are gaining attention as a potential alternative feed source due to their abundant availability and low cost of production [7]. They are a nutritious source of protein, fat, vitamins, fiber, and minerals [8]. They can be reared on organic waste which can reduce waste management costs and will also reduce environmental pollution [9, 10]. Poultry manure produces a substantial amount of biowaste [11], and utilizing this waste to rear insects offers a cost-effective and sustainable alternative for preparing insect-based feeds [12].

Maggots are the larval forms of common flies (*Musca domestica*), and possess a rich nutritive profile with high protein and ether content [13, 14]. Yellow mealworms are the larval forms of the *Tenebrio molitor* beetle and are recognized as an excellent protein source for poultry [9, 13, 15]. Both maggots and mealworms can be reared with minimal resources, utilizing various food industry by-products, thereby reducing food waste and promoting sustainable practices [12, 16, 17]. These larval forms have high protein content and are richer in essential amino acids compared to other concentrates used in poultry, such as soybean and groundnut cake [13, 15]. The use of mealworm meal (MW) and housefly maggot meal (HFM) is common worldwide, but Pakistan’s poultry industry primarily relies on costly conventional feeds. This study aims to identify affordable alternatives to conventional soybean and fish meals. The research is significant as it has the potential to address the challenges of rising feed costs faced by the Pakistani poultry industry. By evaluating the nutrient content of various dietary sources, we aim to provide evidence-based recommendations for enhancing the efficiency and sustainability of poultry nutrition.

## Materials and Methods

The mealworms larvae and maggots were obtained from department of poultry science agriculture university Peshawar where they were reared on wheat barn and poultry waste respectively. Fresh larvae were harvested, and transferred to lab for further analysis. One kg of both dry fish meal and soybean meal were procured from the Indus poultry feed mill, Nizampur, Nowshera.

### Handling of Samples

All samples were kept in sealed air tight plastic bags at -4°C prior to lab analyses. Samples were dried in oven at 70°C and grinded to pass through a sieve size of 0.5 mm. The finely crushed samples were shifted to the poultry nutrition laboratories of the poultry science department and Pakistan Council of Scientific and Industrial Research (PCSIR) labs for further analysis. The analysis for each ingredient was performed in three repetitions.

### Proximate Composition

The samples were analyzed for dry matter, ash content, crude protein, crude fat and crude fiber, following the methods of Association of Official Analytical Chemists [18]. All the experiments were performed in animal nutrition laboratory.

### Determination of Dry Matter and Ash Content

For the estimation of dry matter (DM) and ash content, 2g of each feed sample was measured and placed into clean and pre-weighed crucibles. The crucibles were placed in oven for 18 hours at 100°C. After drying process, the samples were allowed to cool in a desiccator for 30 minutes before being re-weighed. The DM percentage was calculated using the following formula:

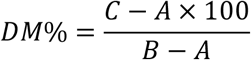

Where:

AA = Weight of the empty crucible (grams)

BB = Weight of crucible + Sample (pre-drying) (grams)

CC = Weight of crucible + Sample (post-drying) (grams)

For the determination of ash content, the samples were incinerated in a muffle furnace at 550°C for 6 hours. After incineration, the samples were cooled in a desiccator and re-weighed. The ash content was calculated by the formula given below:

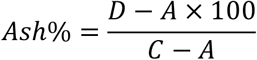

Where:

A =Weight of the empty crucible (grams)

C = Weight of crucible + Sample (post-drying) (grams)

D = Weight of crucible + Ash (grams)

The organic matter (OM) in the feeds was calculated after subtracting the ash percentage from 100%:

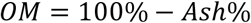

### Determination of Crude Protein

We used Kjeldhal method for crude protein analysis. In each experiment, 0.5g of respective feed sample was weighed and placed in a digestion flask; 4g of digestion mixture (K_2_SO_4_:CuSO_4_:7:1) and 10ml of conc. H_2_SO_4_ was added; the contents of the flask were thoroughly mixed. The mixture was heated under a fume hood until complete digestion, resulting in a greenish-blue color, after which, the flask was allowed to cool. The digest was transferred to a 100ml volumetric flask and volume was made up to the mark using distilled water.

The distillation of the digest was carried out using Markam Still distillation apparatus. 10ml of the digest was added in the distillation tube with the help of funnel; 10ml of 40% NaOH solution was gradually added in the same way. Distillation continued for 10min, producing NH_3_ which was collected as NH_4_OH in a conical flask containing 20ml of 4% boric acid solution and few drops of modified methyl red indicator. The color change from pink to yellowish indicated the presence of NH_4_OH. The distillate was titrated against standard 0.1N sulfuric acid until pink color appeared. A blank sample was also run following similar steps.

The formula for calculating nitrogen percentage was applied as follows:

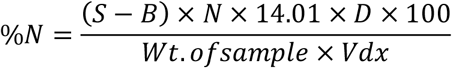

Where:

S = Sample titration reading

B = Blank titration reading

N = Normality of sulfuric acid

D = Dilution of the sample after digestion

V = Volume taken for titration

0.014 is the milli-equivalent weight of nitrogen.

Crude protein was calculated using the following formula for each feed;

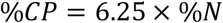

### Determination of Crude Fat

Crude fat was determined by ether extract method using Soxhlet apparatus. One gram of the dried sample was weighed, wrapped in filter paper and placed in thimble. The Soxhlet apparatus was assembled with the extraction tube and receiving beaker. The receiving beaker was pre-weighed to establish a baseline weight. About 100-150ml ether was added to the receiving beaker connected to the Soxhlet apparatus. The thimble containing the sample was fitted into the extraction tube. The extraction continued for up to 4-6 siphoning, extracting fat-soluble components from the sample using the solvent (ether). After extraction process, ether was allowed to evaporate on water bath. Upon complete drying, the receiving beaker was re-weighed to determine the weight of the extracted ether and fat components.

The percentage of ether extract was calculated as follow:

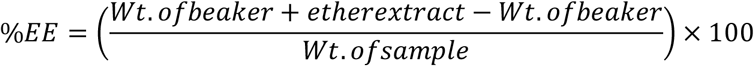

### Determination of Crude Fiber

Two grams of each sample was separately transferred to a beaker and 100ml of 0.1M HCl was added. The beaker which was covered with a condenser, was intensively heated for 5 minutes on a heater, the heat was then adjusted to maintain uniform boiling for 30 minutes. After cooling, 50ml of NaOH solution was added and the mixture was boiled again for 35 minutes. Five minutes before the boiling endpoint, 0.5g Na2EDTA was added. The solution was filtered into pre-weighed crucible, attached to filtration apparatus. Post-filtration, the beaker was rinsed twice with hot water, followed by rinsing with 50ml of 0.3M HCl solution and afterwards again with hot water to remove residual acidity. The beaker was rinsed again with 50ml acetone twice. The crucible was dried in an oven at 100°C overnight. Crucible was allowed to cool, and re-weighed. Crucible was again placed in muffle furnace at 550°C for two hours, cooled and re-weighed.

The percentage of crude fiber was determined by the following formula:

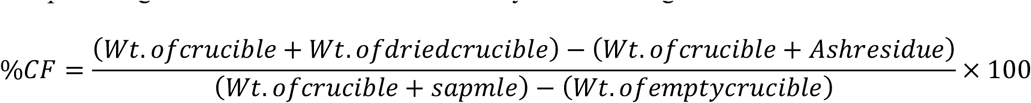

### Amino acid Analysis

We used modified PICO TAG method for amino acids analysis. The analysis was done at the PCSIR lab Peshawar, using high performance liquid chromatography (HPLC) based on the methodologies of [19]. For the experiment, 500mg of each sample was finely ground and hydrolyzed by adding 4.83g barium hydroxide and 5ml of boiling water. The mixture was evacuated and heated at 120°C for 8 hours. After hydrolysis, the pH was adjusted to 3 using HCl, and the solution was diluted to 25ml with HPLC-grade distilled water. Subsequently, 1ml of the sample was vacuum-dried using a flash evaporator and finally dissolved in citrate buffer (0.1M; pH 2.2). Acid hydrolysis was done with 6N HCl at 110°C for 18-22 hours in evacuated and sealed tubes. The hydrolysate was filtered and diluted to 250ml. Another 1ml sample was vacuum evaporated at 40°C until dryness and the content was dissolved in citrate buffer (0.1M: pH 2.2). For HPLC analysis, 20ml of these derivatives were injected into the HPLC system equipped with a Shimadzu HPLC detector LC – 10A, with a variable wavelength monitor set at 350-450nm. The resolution of amino acid derivatives was achieved through a binary gradient system involving solvents A and B. Solvent A consisted of 58.8g of sodium citrate containing 0.2N sodium (pH 3.2), 210ml of 99.55 ethanol, and 50ml (60%) perchloric acid, while Solvent B consisted of 58.5g of sodium citrate containing 0.6N sodium (pH 10), 12.4g boric acid, and 30ml of 4N NaOH solution. Solvents were delivered to the column at a flow rate of 4ml/min for 7 to 10 minutes.

### Determination of Minerals Content

The minerals calcium, phosphorus, magnesium, zinc, and iron contents were determined by atomic mass spectrometry according to the method proposed by [20]. For experiment, samples were dried at 65°C for 48 hours and finely crushed to pass through 0.5mm sieve. Approximately, 300mg of samples were weighed into 20ml calibrated digestion tubes. Care was taken to avoid any adherence of particles to the tube’s sides. In a fume cupboard, 6ml of concentrated nitric acid (70%) was added to each digestion tube and allowed to sit overnight for sample pre-digestion. Subsequently, 2ml of concentrated perchloric acid (70%) was added using calibrated dispenser all while twirling to ensure thorough washing of all samples with acid below the 20ml graduation mark on the tube. All tubes were placed on an aluminum digestion block (AIM 600, Equilab, Canada), initially at a set temperature of 85°C for 45 minutes and then at 110°C for 120 minutes. The temperature of the digestion block was further increased to 160°C until all nitric acid boiled off, leaving only perchloric acid behind. Upon completion of the digestion, all tubes were removed and allowed to cool. Deionized water was added up to the graduation mark in each digestion tube. Samples were vortexed, allowed to settle down, while supernatant was transferred to 25ml PP tubes with lids.

An atomic absorption spectrophotometer was used to determine the contents of calcium (Ca), phosphorus (P), magnesium (Mg), zinc (Zn), and iron (Fe) in each tube against the standard. The readings on the atomic absorption spectrophotometer showed the concentration for each mineral in the samples in terms of weight per unit volume.

The weight of each mineral in the sample was calculated using the equation:

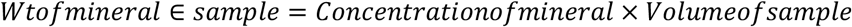

The percentage content of each mineral was calculated using the equation:

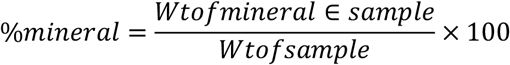

Metabolizable energy calculation

Metabolizable energy (ME) of the feeds were calculated using [21] equation:

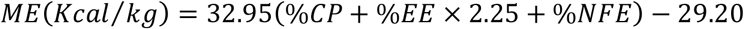

- The constant 32.95 represents the estimated energy density (kcal/kg) of the feed
- The constant -29.20 is subtracted as an adjustment factor to account for energy losses and the presence of anti-nutritional factors

Nitrogen free extract (NFE) was calculated using the following equation:

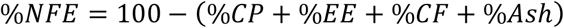

### Statistical Analysis

All data were expressed as mean ± standard error (SE) of three replicates. Nutrient composition data were analyzed using one-way analysis of variance (ANOVA) to determine significant differences among the feed ingredients. All statistical analyses were conducted using SPSS software (version 26.0; IBM Corp., Armonk, NY, USA).

## Results

### Proximate Analysis

We observed no significant differences in the dry matter content among fish meal, mealworms, and HF maggot (p > 0.05). The mean crude protein values for maggot (57.8) and fish meal (58.5) were comparable, but significantly differed from the CP values recorded for mealworms (51.02) and soybean meal (45.8) (p < 0.05). The highest ether extract value was recorded for mealworms (27.3%) and maggot (26.9%), followed by comparatively lower value for fish meal (21.5%) and least was recorded for soybean meal (2.9%). Crude fiber content was recorded highest for mealworms and soybean meal while housefly and fish meal had low CF content. No significant difference was found between Ash content of mealworms (5.7%) and maggots (5.9%) (p > 0.05), and between soymeal (6.2%) and fish meal (7.1%) (Table 1). The nitrogen free extract (NFE) was found highest in soybean meal (39.5%) indicating high carbohydrate content. The NFE contents in fish meal and mealworms were 11.2% and 9.7% respectively. The least NFE value was recorded for HFM (6.9%) (Table 2).

**Table 1.** Proximate analysis of dried mealworm, housefly maggot, fish meal and soybean meal (mg/kg)

|  | Dry matter % | Crude protein % | Ether extract % | Crude fiber % | Ash% |
| --- | --- | --- | --- | --- | --- |
| | Mean ± SE | Mean ± SE | Mean ± SE | Mean ± SE | Mean ± SE |
| Mealworms | 94.1 ± 0.32 | 51.02 ± 0.18 | 27.3 ± 0.52 | 6.2 ± 0.21 | 5.7 ± 0.23 |
| Housefly<br>maggots | 94.2 ± 0.35 | 57.8 ± 0.28 | 26.9 ± 0.28 | 3.1 ± 0.17 | 5.9 ± 0.2 |
| Fish meal | 93.9 ± 0.23 | 58.5 ± 0.26 | 21.5 ± 0.27 | 1.7 ± 0.14 | 7.1 ± 0.16 |
| Soymeal | 91.8 ± 0.32 | 45.8 ± 0.21 | 2.9 ± 0.14 | 5.6 ± 0.21 | 6.2 ± 0.15 |

The number after ± represents standard error value

**Table 2.**
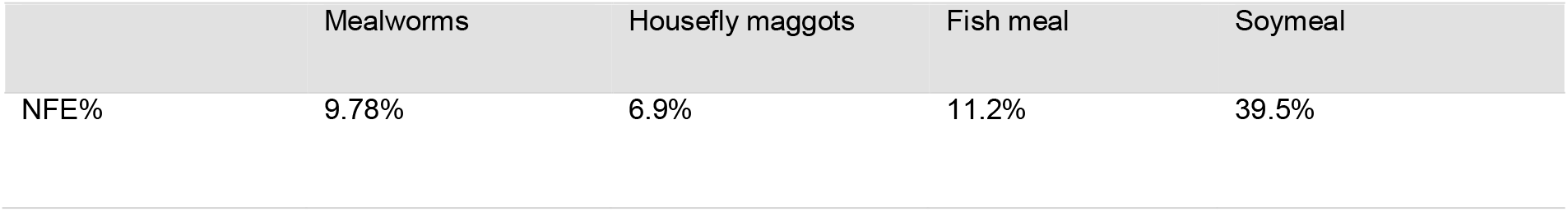
Nitrogen free extract values of dried mealworms, housefly maggots, fish meal and soymeal.

### Mineral Content

The highest calcium content was found in mealworms (5.23 g/kg) followed by HFM (5.13 g/kg) (p > 0.05). These values were higher than the values recorded for fish meal (3.57 g/kg) and soybean (3.7 g/kg). The highest phosphorus content was found in maggot (16.8 g/kg). The phosphorus content values in mealworms (4.87 g/kg), fish meal (3.18 g/kg), and soybean meal (5.43 g/kg) were relatively close to each other. The highest value for magnesium content was observed in fish meal (23.2 mg/kg), followed by maggot at 4.07 mg/kg value. The highest iron content was recorded in soybean meal at 116.5 mg/kg, followed by fish meal at 20.9 mg/kg. The highest zinc content was found in HFM (98.5 mg/kg) while least value was recorded for soybean meal (9.8 mg/kg) (Table 3).

**Table 3.** Mineral content of dried mealworm, maggot, fish meal and soybean meal.

| Minerals | Mealworms | Housefly maggots | Fish meal | Soybean meal |
| --- | --- | --- | --- | --- |
| | Mean ± SE | Mean ± SE | Mean ± SE | Mean ± SE |
| Calcium (g/kg) | 5.23 ± 0.2 | 5.13 ± 0.17 | 3.57 ± 0.14 | 3.7 ± 0.2 |
| Phosphorus (g/kg) | 4.87 ± 0.13 | 16.8 ± 0.28 | 3.18 ± 0.09 | 5.43 ± 0.26 |
| Magnesium (mg/kg) | 2.66 ± 0.28 | 4.07 ± 0.14 | 23.2 ± 1.04 | 2.53 ± 0.12 |
| Iron (mg/kg) | 4.53 ± 0.14 | 1.83 ± 0.20 | 20.9 ± 0.14 | 116.5 ± 1.72 |
| Zinc (mg/kg) | 27.5 ± 0.2 | 98.5 ± 1.04 | 72.1 ± 0.71 | 9.8 ± 0.16 |

The number after ± represents standard error value

### Amino acid Profile

The most limiting amino acids lysine and methionine contents were found highest in housefly maggot (6.53:2.27). The lysine and methionine values were also found high in mealworms (5.51:2.3) and fish meal (5.57:2.01) with only a slight difference between their values which indicates comparable status. The least lysine and methionine values were recorded for soybean meal (2.85:0.71). The highest leucine and isoleucine values were recorded for HFM (6.71:4.1), followed by fish meal (6.1:3.82) and mealworms (5.9:3.1). The minimal differences observed among the values for housefly maggots, mealworms and fish meal indicates comparable amino acid profiles of these feed ingredients (Table 4).

**Table 4.** Amino acid profile (g/100g protein) of dried mealworm, housefly maggot, fish meal and soybean meal.

| Amino acid | Mealworms | Housefly maggots | Fish meal | Soybean meal |
| --- | --- | --- | --- | --- |
| Arginine | 2.31 | 6.17 | 4.1 | 2.58 |
| Cystine | 4.3 | 1.05 | 0.88 | 0.92 |
| Histidine | 1.15 | 3.4 | 1.13 | 1.05 |
| Isoleucine | 3.1 | 4.1 | 3.82 | 2.23 |
| Leucine | 5.9 | 6.71 | 6.1 | 3.4 |
| Lysine | 5.51 | 6.53 | 5.57 | 2.85 |
| Methionine | 2.3 | 2.27 | 2.01 | 0.71 |
| Phenylalanine | 1.1 | 4.38 | 3.1 | 2.21 |
| Threonine | 0.9 | 3.34 | 3.14 | 1.53 |
| Tryptophan | 0.81 | 0.43 | 0.95 | 0.91 |
| Valine | 2.9 | 4.53 | 3.38 | 2.87 |

### Metabolizable Energy (ME) kcal/kg

The highest metabolizable energy was recorded for housefly maggots at 4,074 kcal/kg value, an indication that housefly maggots provide the most energy per kilogram compared to the other feed ingredients. The ME value recorded for mealworms was 3,988 kcal/kg which is slightly higher than the fish meal value (3,824 kcal/kg), showing their comparable status. The soybean meal had lowest ME value (2,995 kcal/kg), indicating lower energy density compared to other evaluated feed sources in this study (Table 5).

**Table 5.** Metabolizable energy (ME) contents of the dried mealworms, housefly maggots, fish meal and soymeal.

|  | Metabolizable Energy (ME) |
| --- | --- |
| Mealworms | 3,988 kcal/kg |
| Housefly maggots | 4,074 kcal/kg |
| Fish meal | 3,824 kcal/kg |
| Soybean meal | 2,995 kcal/kg |

## Discussion

Insects are considered an excellent source of nutrients for poultry due to high protein content, balanced amino acid profile and energy levels [22]. Insect rearing enhances food and feed security due to easy growth, high feed conversion efficiency, and utilization of bio-waste [23]. The mealworm (MW) are the larval forms of darkling beetles (*Tenebrio molitor*), and contain high concentrations of protein, fat, amino acids, and minerals [5]. Several studies have investigated the impact of incorporating mealworms into poultry feed [24]. Maggots, the larval forms of common flies (*M. domestica*), also provide essential protein and other nutrients for poultry [13]. Housefly maggots, known for their ability to grow on various substrates, can convert waste into valuable protein and fat-rich biomass [23, 25].

Poultry feed traditionally focuses on providing energy, but improving nutrient balance, like protein and amino acids greatly benefits chick development [26]. Protein is essential for carcass development, egg production, feather formation and other vital physiological functions [26, 27, 28].

Increased crude protein content in feeds results in improvement in egg size and weight while a lower CP content will affect development of carcass and eggs production [26]. In our study, no significant difference (*p* > 0.05) was observed in crude protein (CP) content between housefly maggots (57.8%) and fish meal (58.5%), which had the highest recorded CP value, indicating that HFM is comparable to fish meal in terms of protein content. This is in accordance with the previous studies where maggots were observed to possess high protein content (30–64%) and essential amino acids (EAA), exceeding the protein quality of soybeans and competing with fish meal [15, 25]. Our recorded CP content for housefly maggots aligns with the CP values reported previously; 55.6% [29], 63.99% [30], 62.98% [31], 50% [32], 55.4 [33], 42.70, 48.4 [34]. The ether extract (EE) content in maggots was 26.9%, slightly lower than the highest value recorded for mealworms (27.3%) (*p* > 0.05). Our observed value of EE for HFM falls within the range of previous studies; 28% [15], 27.9% [29], 24.31% [30]. Variations in nutrient content across studies are likely due to differences in pupal age, rearing substrates, and drying methods [25, 35]. In the mealworms, we recorded a CP content of 51.02%, which is slightly lower than that of maggots and fish meal but significantly higher than the CP content in soybean meal. Mealworms have a usual CP range of 27 to 54%, which is considered optimal and makes them a good source of protein for poultry diets [15, 36]. Our recorded CP value for mealworms is consistent with reports of 46.44% [37], 51.93% [24], 52.89% [31], 53% [32]. We recorded the highest ether extract value for mealworms at 27.3%. This value is consistent with previous reports of 30.05 [31], 29.6 [38]. The crude fat content in mealworms has been observed to range widely from 4% to 34% [15, 36]. This broad range can be attributed to factors like variations in mealworms diet, and environmental conditions.

We recorded the highest metabolizable energy (ME) in maggots at 4,074 kcal/kg, followed by mealworms at 3,988 kcal/kg. The high ME value for maggots is consistent with the previously reported value of 3,955 kcal/kg [25]. The high ME values of HFM and mealworms suggested the high protein and fat content of the insect-based sources compared to traditional feeds. The ME value for fish meal was also high at 3,824 kcal/kg, only slightly lower than that of mealworms, indicating comparable quality.

The ash component of the feed describes the inorganic content of the feed, mainly minerals, which are required in specific amounts for stronger bone, blood clotting and egg shell formations (Jacquie, 2018). In the current study, no significant differences (*p* > 0.05) were observed in the dry matter and ash contents of mealworms (94.1%, 5.7%) and HFM (94.2%, 5.9%). The DM and ash contents in HFM are consistent with the previous reports: 92.7% DM and 6.23% ash [33], 92.3% DM and 4.20% ash [39], 93.9% DM and 14.5% ash [34]. In the minerals profile, the highest calcium content was found in mealworms (5.23%), followed closely by maggots (5.13%). The calcium content in maggots and mealworms was higher compared to the calcium values observed in fish and soybean meal. The calcium content for HFM was consistent with the 4.9 g/kg reported in a previous study [25]. The calcium content reported in our study for mealworms was comparatively higher than previously reported values: 0.25 [31], 0.05 [38], 0.43 [37]. This suggests that insect meals may be deficient in calcium, but its levels can be enhanced by manipulation of the substrate on which insects are reared [23]. The phosphorus content for maggots was 16.8%, the highest in our study, and is consistent with previous report of 16% [25]. The phosphorus content in mealworms was 4.87%. Wide range differences have been observed among the observed phosphorus contents in various studies, such as 7.06 [37], 0.72 [38], 0.74 [31].

The amino acids leucine and isoleucine are often implicated in amino acid antagonism and need to be considered in poultry feed formulation [27, 40]. In the current study, essential amino acids (EAAs) isoleucine, leucine, lysine, valine contents were highest in maggots with 4.1, 6.71, 6.53 and 4.53 values respectively. The EAA amino acid methionine content was found highest in mealworms (2.3) followed by maggot (2.27). These values are in accordance with the previous reports where leucine, isoleucine, lysine and methionine contents in maggots were found higher than fish and soybean meal [25]. The tryptophan content in mealworms was 0.81 while in HFM its value was 0.43 which is least among the evaluated diets in our study. Few studies have shown poor performance of broilers with low level of tryptophan in maggot meal, while mealworms have been found to have higher tryptophan content comparatively [24]. In the mealworms, our recorded values for isoleucine, leucine and lysine were 3.1, 5.9 and 5.51 respectively. These values are higher than those for soybean meal, which is consistent with reports that lysine, methionine, tryptophan, and threonine contents in dried mealworms are comparable to soybean meal [31]. The amino acids profiles of mealworms, fish and soybean meals have been found to be closely associated [31, 41].

Many researchers encourage the partial replacement of soybean and fish meal with maggot meal and worm meal in poultry rations [23, 25]. Surveys have shown the willingness of poultry farmers to use insect based meals as alternative protein source in poultry diets [42]. The palatability of these feeds is good, and they can partially replace soybean or fish meal depending on the species.

## Conclusion

The comparative analysis of the evaluated feeds showed that housefly maggots and mealworms have high levels of crude protein, ether extract, and essential amino acids compared to soybean meal, and are comparable in quality to fish meal. The mineral profiles of both insect meals were also similar to those of fish meal and superior to soybean meal. Energy content analysis revealed that housefly maggots provided the highest metabolizable energy, offering more energy per kilogram than the other feed ingredients evaluated in this study. These results are promising, highlighting the great potential of insect-based meals as sustainable alternatives to conventional poultry feed ingredients. Author Contributions

All authors contributed to the study conception and design. GN and SK conceived and designed the study. GN performed all the experimental work. SBR and AS assisted with data analysis, while RK and IU contributed to manuscript review. GN wrote the initial draft, which was finalized under the supervision of SBR and SK. All authors read and approved the final manuscript.

## Statements and Declarations

Consent to participate

There are no human participants in this article and informed consent is not required.

## Declaration of Conflict of Interests

The authors declare no conflict of interest.

## Acknowledgments

We thank the Pakistan Science Foundation for their support of this project.

## Funding Statement

The author(s) received no financial support for the research, authorship, and/or publication of this article.

## Notes

### Competing Interest Statement

The authors have declared no competing interest.

